# Neural Innervation Invigorates Yolk Sac Biological Functions beyond Nutrient Reservoir during Zebrafish Embryo Development

**DOI:** 10.64898/2026.06.06.730572

**Authors:** Zhengduo Wang, Li Tian, Bo Li

## Abstract

The zebrafish yolk sac (YS) is traditionally viewed as a nutrient reservoir. By reconstructing the complete progression of embryonic neural development via live-cell imaging, we previously observed that axonal projections extending from brain cranial neurons come to cover the yolk sac surface, hinting at its uncharacterized functional roles beyond nutrient storage. Using transgenic lines and long-term live imaging, we characterized a dynamic neuro-vascular-metabolic interface on the YS surface. We observed that peripheral neural networks expand radially and mature through hierarchical integration, sharing the same structural and dynamical features as those in the brain and spinal cord. Unexpectedly, YS surface cells harbor transient calcium flashes, reflecting primitive intercellular communication in response to neural innervation. Furthermore, we characterized activity-dependent neuronal pruning and stress-induced lipid droplet crystallization as indicators of developmental refinement and homeostatic collapse, respectively. Finally, we identified directional blood flow occurring before the formation of endothelial tubes, indicating a pre-vascular transport mechanism. These findings demonstrate that neural innervation enables the YS to serve as a coordinated developmental hub, facilitating complex crosstalk between neural, vascular, and metabolic systems during early vertebrate embryogenesis.

**Significance Statement:** Traditional paradigms view the embryonic yolk sac simply as a nutrient reservoir. Using advanced live-cell imaging in zebrafish, we uncover that the yolk sac can be innervated by brain-expanded axonal networks during development, invigorating its biological functions, including neural signal, vascular flow, and metabolic activity. Rather than maintaining local neuronal bodies, these long-distance nerve extensions map out an early topological scaffold that may guide future internal organ innervation during yolk resorption. Furthermore, we capture pre-vascular blood transport and stress-induced metabolic shifts on the yolk sac surface. These findings redefine the yolk sac as a neural-controlled functional hub before organs form.

## Introduction

The zebrafish (Danio rerio), with its unique advantages of embryonic transparency, external development, high genetic tractability, and rapid development, has become a core model organism for dissecting the mechanisms of early vertebrate development (Lessman, 2011; Chia et al., 2022). In our previous work, we established a live-cell imaging platform for zebrafish embryos and captured and characterized the entire process of neurogenesis and its guiding role for embryogenesis (Wang et al., 2026). During imaging, we observed that axonal networks extending from the brain covered the yolk sac (YS), suggesting that the yolk sac may have functions beyond nutrient storage. To further elucidate these non-nutritive roles of the yolk sac in development, we used our established imaging platform to perform an in-depth investigation.

Traditionally viewed primarily as a passive nutrient reservoir for early embryogenesis via biochemical regulation (Quinlivan and Farber, 2017; Chen et al., 2024), the zebrafish YS also acts as a signaling center. During the first 5 dpf, yolk proteins are hydrolyzed within the I-YSL—enriched with lysosomes and ER—and delivered to embryonic cells via transport or diffusion (Carvalho and Heisenberg, 2010; Quinlivan and Farber, 2017; Furukawa et al., 2024; Sun et al., 2026). Early, the external YSL modulates Nodal/Wnt signals to guide gastrulation (Feldman et al., 1998; Bischof and Driever, 2004). Later, YSL-specific transcription factors and YS-secreted cytokines regulate ECM assembly, cardiomyocyte migration, and immune development (Sakaguchi et al., 2006; Kawahara et al., 2009; Iturri et al., 2021; Diez-Pinel et al., 2026). Cellular-level studies reveal the cellular infrastructures of YS that support its biological functions as a nutrient reservoir and transient biochemical dispenser (Fukazawa et al., 2010). Between 24 and 48 hpf, the YS surface assembles into a well-organized, sandwich-like tri-layer architecture (Fukazawa et al., 2010): an outer periderm barrier sealed by tight junctions (Kimmel et al., 1990; Kiener et al., 2008; Fukazawa et al., 2010), an intermediate EBL secreting collagen for the basement membrane (Bader et al., 2009; Takeuchi et al., 2015; Sampedro et al., 2018), and the deep YSL enwrapping the internal yolk mass (Carvalho and Heisenberg, 2010).

Despite these structural insights, current paradigms have not yet considered whether the yolk sac participates in neural development and regulation in a biophysical or systemic manner. Our previous work reveals that the YS surface is innervated by axons projecting directly from cranial neurons in the brain, with their somata strictly remaining in the central nervous system (Wang et al., 2026). This provide a hint that rather than being a localized auxiliary organ, the YS may function as an integrated structural and physiological hub that coordinates whole-body morphogenesis. However, the precise mechanisms underlying this neural-YS interplay remain poorly understood. Whether and how the nervous system functionally dictates the yolk sac, and how the complex vasculature is biophysically established on its surface, remain critical unanswered questions.

To elucidate these multidimensional regulatory mechanisms, this study employed multiple specific transgenic zebrafish lines (Tg(elavl3:EGFP), Tg(elavl3:GCaMP6f) and Tg(fli-1:mCherry)) combined with high-resolution long-term live imaging to systematically characterize the topological evolution of the peripheral neural network on the YS surface, the collective calcium dynamics of surface cell populations, the non-vascular-dependent early fluid dynamics, and the reorganization of lipid metabolism under stress during early zebrafish development. Our results indicate that the YS is not only nutrient-bearing organ as traditionally perceived, but also serves as functional developmental hub exhibiting high spatiotemporal coordination. On the giant curved surface constructed by its microenvironment, there exist profound multi-field reciprocal interactions and spatiotemporal synergy among the nervous system, the immature vascular flow field, and the lipid metabolic network. These findings provide a novel perspective for understanding multisystem crosstalk and the regulatory principles of the extra-embryonic microenvironment on overall development.

## Materials and Methods

### Construction and maintenance of transgenetic zebrafish lines

Zebrafish strains Tg(elavl3:EGFP) and Tg(elavl3:GCaMP6f) were acquired from the China Zebrafish Resource Center (http://www.zfish.cn/), and the Tg(fli-1:mCherry) strain was purchased from Nanjing Ezerinka Biotechnology Co., Ltd. (http://www.ezerinka.com/). The hybrid lines (elavl3:EGFP; fli-1:mCherry) were generated in-house through crossbreeding and selective rearing. All zebrafish were maintained in a recirculating aquaculture system (ZF-104, Nanjing Ezerinka Biotechnology Co., Ltd.).

The system temperature was set to 28□°C, with continuous operation of UV sterilization and oxygen supply. A 5% sea salt solution and 5% sodium bicarbonate solution were used to regulate conductivity and ion balance, maintaining system conductivity between 500–800□μS/cm and pH between 7.0–8.0. Ambient temperature was controlled at 28□°C using air conditioning, and a 12□h light/12□h dark photoperiod was applied. Approximately one-third of the system water was replaced daily to keep total ammonia nitrogen below 0.02□mg/L.

To facilitate subsequent spawning, male and female zebrafish were housed separately. Males were distinguished by their slender body shape, flat abdomen, and lemon-yellow coloration, whereas females exhibited a fuller body, rounded abdomen, and silvery-gray pigmentation, allowing reliable visual sex identification. Adult zebrafish were fed live Artemia nauplii daily. The hatching procedure was as follows: 5□g of Artemia cysts and 5□g of sea salt were added to 1□L of pure water. The mixture was aerated under illumination for 24□h, after which aeration was stopped to allow eggshells to separate. The hatched nauplii settled at the bottom and were collected for feeding. Fish were fed 1-2 times per day, with the amount adjusted to ensure consumption within 15-20 minutes.

### Sample preparation for *in vivo* zebrafish imaging

On the day before imaging, female and male zebrafish of different transgenic strains were placed in spawning tanks separated by a divider. The tanks were filled with a 1:1 mixture of fresh water and system water, covered, and maintained overnight. On the following morning, after the onset of illumination, the divider was removed to allow direct interaction between males and females. Males began nudging the abdominal region of the females, which subsequently initiated spawning. Approximately 30 minutes later, both females and males were returned to the main rearing system. The fertilized eggs were collected from the spawning tanks into 10 cm Petri dishes and maintained in specialized zebrafish embryo medium at 28□°C.

Embryo imaging typically commenced around 24 hours post-fertilization (hpf). Prior to imaging, embryos were screened using a spinning disk confocal microscope, and only those exhibiting clear fluorescence signals were selected for further processing. The chorion was then removed from each embryo to improve imaging quality. Although the chorion serves as a protective barrier during early development, it limits imaging depth and introduces motion artifacts as embryos become more motile inside the membrane. For dechorionation, 24-hpf embryos were transferred to a Petri dish with a small amount of water and placed under a stereomicroscope. A small incision was made in the chorion using a corneal scissors, and the embryo was gently extruded with fine forceps.

The dechorionated embryos were then transferred to a confocal imaging dish. Low-melting-point agarose (0.12□g) was dissolved in 10□mL of ddH2O by microwave heating to prepare a 1.2% solution. After cooling to approximately 35□°C, the agarose solution was dispensed over the embryos. The position and orientation of each embryo were adjusted with forceps before the agarose solidified. For imaging at earlier developmental stages, such as 12 hours post-fertilization (hpf), the chorion was not removed due to the structural fragility of the embryos at this period, as mechanical manipulation could readily cause damage. Instead, the embryos within their intact chorions were directly embedded in low-melting-point agarose for immobilization during imaging.

### Live cell imaging

Live imaging of zebrafish embryos was performed using a Spin SR10 spinning disk confocal microscope (Evident Olympus) equipped with an Oko-lab incubation chamber (H301-MINI). The system maintained a constant temperature of 28□°C throughout the experiments. Embryos were mounted in confocal dishes (Nest, 801001) and imaged with a typical laser intensity set at 20% and exposure time at 200□ms. For fluorescence excitation, the 488□nm laser line was used to visualize pan-neuronal labels in the transgenic lines (elavl3:EGFP, elavl3:GCaMP6f), while the 561□nm laser was employed for imaging the (fli-1:mCherry) line.

To enhance experimental throughput, approximately 20 embryos were placed in each confocal dish. The microscope’s positional memory function was utilized to record individual embryo locations, enabling simultaneous time-lapse acquisition from multiple specimens. All embryos were imaged using axial Z-stack acquisition mode with the following parameters. Z-stack imaging was performed witha 30x oil objective at 200 μm Z-range and 5-6 μm step size, and a 20x objective at 400 μm Z-range and 10 μm step size, respectively. Time-series imaging was performed at 10-minute intervals over a total duration of 48 hours, generating continuous 3D volumetric data throughout the recording period.

### Data reconstruction and 3D visualization

The reconstruction of imaging stacks from confocal microscopy is conducted using Cell Sens Dimension Desktop 4.2.1 (https://lifescience.evidentscientific.com.cn/en/software/cellsens/). In the top planar view, the Maximum Intensity Projection (MIP) algorithm is adopted to overlay all images in the z-stack into a single image. In the side view, the Marching Cubes (Surface Reconstruction Algorithms) algorithm is used to yield the stereoscopic spheroid. Compared with the range in the xy plane, the length scale in z is relatively small, making the analysis based on xy projection reasonably accurate.

### Analysis of live cell imaging data

Time-lapse imaging data of zebrafish neural network development were analyzed using ImageJ 1.54p. Growth cone dynamics, including migration trajectories and velocity, were quantified with the Manual Tracking plugin. The analysis workflow consisted of the following steps: Time-lapse image stacks were loaded into Fiji, and the Manual Tracking plugin was initialized with appropriate temporal intervals and spatial calibration based on the acquisition settings. Growth cones were manually selected and tracked across consecutive frames using the plugin’s point-and-click interface. The plugin automatically recorded their positional coordinates over time and computed kinematic parameters such as displacement, velocity, and persistence. To assess structural organization within the developing network, angular relationships between neurites were also measured using Fiji’s built-in angle measurement tool. Specific neurite segments and growth cone orientations were manually annotated, and their relative angles were calculated to evaluate network topology and directional guidance. All tracking and angular data were exported for further statistical analysis and graphical representation in GraphPad Prism 8.

## Results

### Topological Expansion and Hierarchical Maturation of Neural Network on the Yolk Sac Surface

To observe the neural development on the YS surface in real time, we used the pan-neuronal transgenic line Tg(elavl3:EGFP), which specifically expresses enhanced green fluorescent protein (EGFP) in mature neurons (Park et al., 2000). We performed long-term live confocal imaging of zebrafish embryos from 24 to 48 hpf, with an imaging interval of 10 min and a typical z-step of 5 μm, over a z-scan range exceeding 200 μm to ensure complete coverage of the three-dimensional structure of the YS surface on one side of the embryo.

Imaging results showed that at 24 hpf, neuronal growth cones began to extend from the embryonic brain and spinal cord toward the YS surface. As development proceeded, growth cones continuously expanded radially onto the YS, forming a dynamically expanding neural network frontier (**Fig.1A-B; Movie S1**). We quantified the movement velocity of growth cones at different developmental time points. The results showed that the growth cone velocity on the yolk sac surface remained relatively stable during the early phase of network expansion and decreased slightly with development; the average growth cone velocity was approximately 0.845 μm/min (**Fig.1C**). The area of the YS covered by the neural network increased approximately linearly over time. Quantification of the uncovered area on the YS surface indicated that it took approximately 10 hours for the growth cones to form a primary neural network covering the YS surface (**Fig.1D**).

**Fig. 1.**
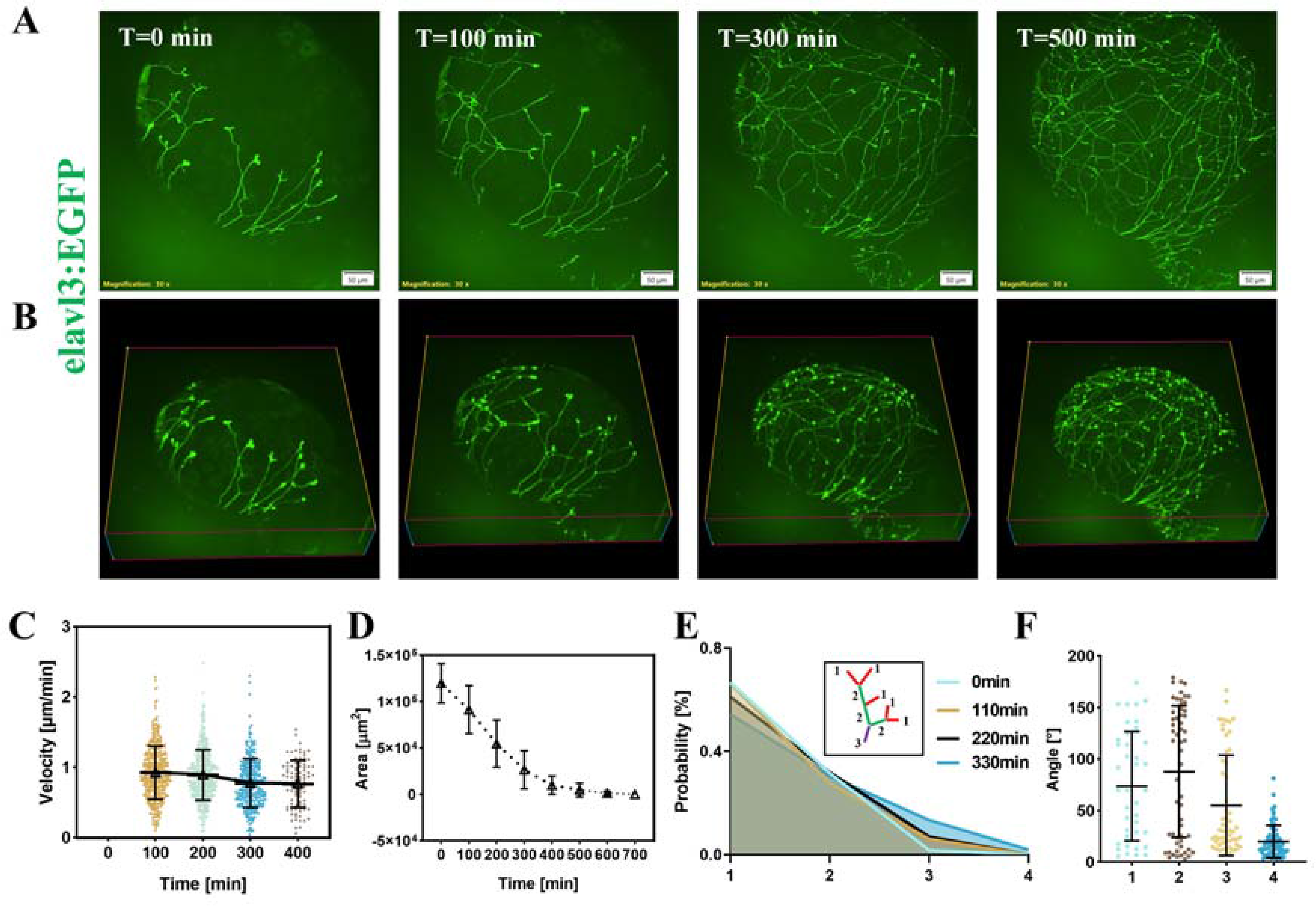
Spatiotemporal expansion and hierarchical topological maturation of neural networks on the zebrafish yolk sac. **(A, B)**, Representative confocal micrographs showing the outgrowth of neural networks on the yolk sac surface of Tg(elavl3:EGFP) zebrafish embryos, starting at 24 hours post-fertilization (hpf). Pan-neuronal signals are labeled with green fluorescence. Time-lapse imaging was performed every 10 min using a 30× silicone immersion objective. Scale bar = 50 µm. (**B**) Corresponding 3D reconstruction of the neural network architecture. (**C**), Quantitative statistics of growth cone migration velocity at distinct developmental stages. (**D**), Line plot showing the uncovered area of the yolk sac surface by neural networks as a function of developmental time. For all statistical plots (n > 3 experiments) was conducted for all data. (**E**), Frequency distribution of neural branches at distinct hierarchical levels during the assembly of yolk sac-surface neural networks. Terminal axons directly extending from growth cones were defined as Level 1. Level 2 branches form via the fusion of two Level 1 branches, and Level 3 branches arise from the convergence of two Level 2 branches. Merging between branches of different hierarchical levels yields a new branch classified according to the higher-order level. (**F**), Statistical quantification of the angular deviation between the overall growth direction of neural networks and axonal orientation at different hierarchical levels. For all statistical plots (n > 3 experiments) was conducted for all data. The error bars are the standard deviation of the data.

To quantitatively characterize the structural features and maturation process of the neural network on the YS surface, we adopted the Strahler analysis method used for stream ordering in hydrology (Zhao et al., 2025), which has been widely applied to study the topology of vascular and neural networks. According to Strahler ordering rules, we classified nerve fibers in the network into different orders: when two fibers of the same order merge, they form a fiber of the next higher order; when fibers of different orders merge, the order remains the higher one. We defined orders 1 and 2 as low-order structures (frontline exploratory branches) and order ≥3 as high-order structures (mature trunks). We calculated the frequency distribution of fibers of different orders at various developmental time points. The results showed that as development progressed, the proportion of low-order fibers gradually decreased while that of high-order fibers increased, indicating that the neural network on the YS surface underwent a maturation process from simplicity to complexity and from dispersion to integration (**Fig.1E**). Further analysis of the spatial orientation of fibers of different orders revealed that the alignment of high-order fibers was highly parallel to the overall growth axis of the neural network, whereas the orientation of low-order fibers exhibited significant randomness (**Fig.1F**). This result suggests that the construction of the YS surface neural network follows a synergistic strategy of “ local exploration--global integration”: low-order fibers are at the growth cone frontier, conducting large-scale spatial exploration through highly random growth to find suitable pathways; those fibers that successfully find targets and are selected for retention merge to form high-order trunk structures responsible for long-distance signal transmission and network integration.

The neural network covering the yolk sac originates from neurons in the zebrafish brain. It is important to emphasize that no neuronal somata are located within the YS itself; rather, all nerve fibers on the YS surface represent long-distance axonal extensions originating from cranial neuronal cell bodies in the brain, a developmental trajectory we previously documented. During early neural development, neuronal somata aggregate to form cell clusters, from which axonal networks extend to cover the brain, spinal cord, and yolk sac surface (Wang et al., 2026). Notably, both the growth cone velocity on the yolk sac surface and the dynamics of neural network formation closely resemble those observed on the brain surface, suggesting potential functional similarities between the two networks. What is the ultimate fate of this YS surface neural network? The traditional view holds that as the YS is absorbed, all structures on the YS surface are degraded and cleared. However, we propose a new hypothesis: as the yolk is gradually consumed, the nerve fibers on the YS surface do not simply disappear but may undergo an involution-like process, entering the embryo interior along with the retracting YS, and transforming into a primitive scaffold for the autonomic nervous system that innervates visceral organs. This hypothesis provides a new perspective for deciphering the origin of the vertebrate peripheral nervous system.

### Calcium Dynamics on the Yolk Sac Surface after Neural Innervation

To investigate the functional activity of the neural network on the YS surface, we used the transgenic zebrafish line Tg(elavl3:GCaMP6f), which expresses the genetically encoded calcium indicator GCaMP6f in pan-neuronal cells (Dunn et al., 2016), enabling real-time monitoring of intracellular calcium concentration changes with high spatiotemporal resolution. We performed calcium imaging on the yolk sac of zebrafish embryos from 24 to 52 hpf, with an imaging interval of approximately 5–10 s, continuously imaging each embryo for 10 min.

Surprisingly, we observed widespread and prominent spontaneous calcium flickers on the YS surface. These calcium flickers manifested as rapid increases in GCaMP6f fluorescence intensity within individual cells, followed by a return to baseline. More importantly, these calcium flickers exhibited clear population burst characteristics: at certain time points, dozens of cells displayed calcium flashes almost simultaneously, forming a calcium wave lasting approximately 10–20 s before synchronous recovery (**Fig.2A-H; Movie S2**). This population burst activity is highly reminiscent of the spontaneous collective neural network activity observed in the zebrafish central nervous system (Avitan et al., 2021; Rosch et al., 2024). Of note, the calcium flickers appear right after the neural innervation by the axons from the brain, suggesting that functional communication is established between the central neural system and the YS, namely the formation of a primitive neural circuit. We further quantified the overall frequency and amplitude of calcium flickers at different developmental stages. The results showed that the flicker frequency gradually decreased with development, from approximately 4 events per 500 s at 36 hpf to about 1 event per 500 s at 52 hpf (**Fig.2I and J**). While the frequency of calcium transients decreased in cells on the yolk sac surface, it significantly increased in neuronal somata within the brain (**Fig.2K**). This decline indicates that early extra-embryonic calcium flashes suggest the biological function of yolk sac surface cells is transient and will disappear eventually.

**Fig. 2.**
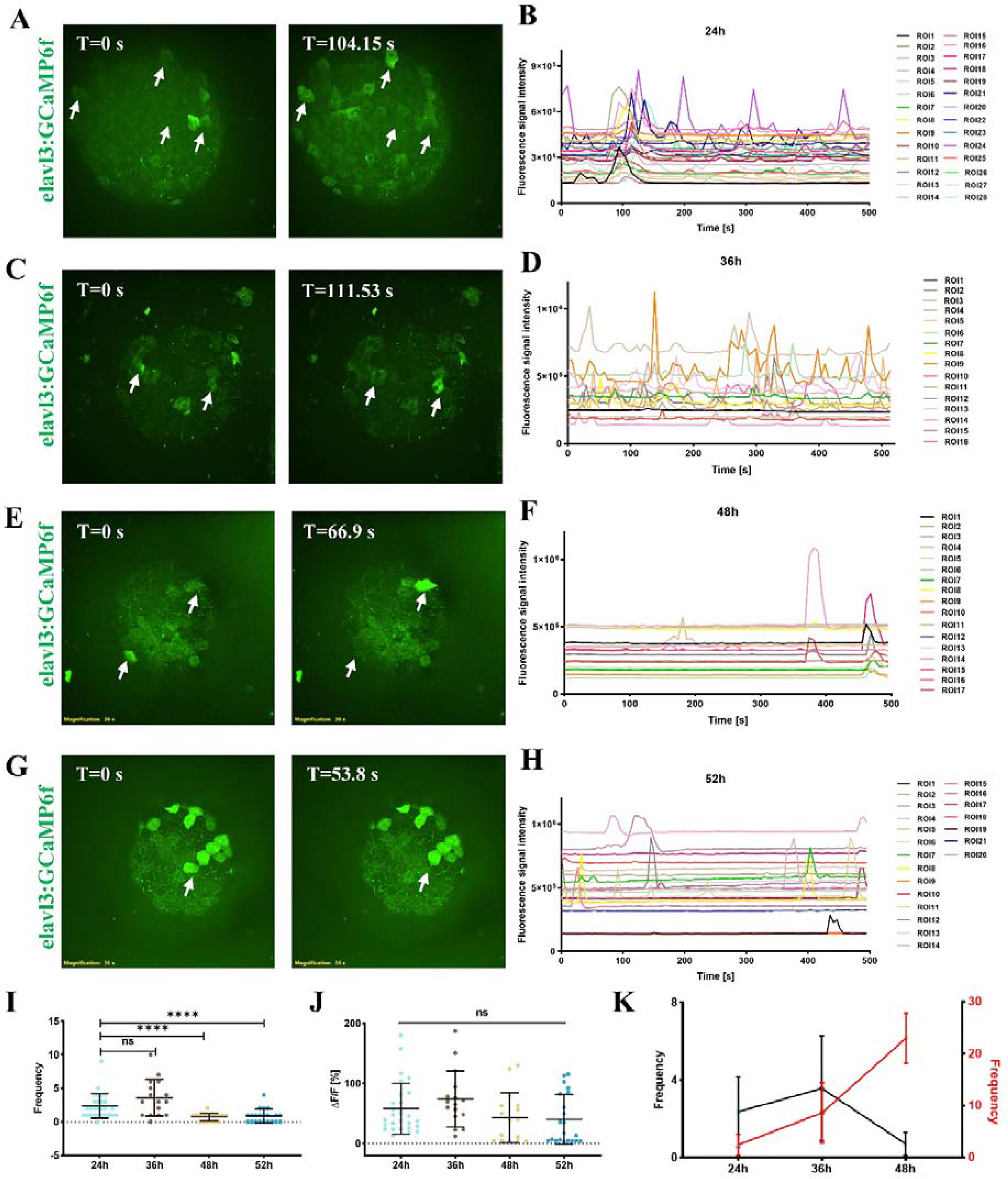
Collective calcium bursting dynamics and developmental changes in elavl3-positive cells on the yolk sac surface. **(A)**, Representative confocal snapshots of spontaneous calcium transients on the yolk sac surface of Tg(elavl3:GCaMP6f) zebrafish embryos at 24 hpf. White arrows indicate individual cells exhibiting prominent calcium flashes. Green fluorescence labels pan-neuronal Ca2+ activity. Imaging was performed using a 30× silicone immersion objective with a frame interval of 10.415 s. Scale bar = 50 μm. (**B**), Raster plot of calcium activity in yolk sac surface cells of zebrafish embryos at 24 hpf over an imaging duration of approximately 500 s; each ROI corresponds to an individual cell. (**C**), Representative confocal snapshots of spontaneous calcium transients on the yolk sac surface of Tg(elavl3:GCaMP6f) zebrafish embryos at 36 hpf. White arrows mark cells with evident calcium flashes. Pan-neuronal Ca2+ signals are visualized by green fluorescence. Imaging was performed with a 30× silicone immersion objective at a frame interval of 5.5765 s. Scale bar = 50 μm. (**D**), Raster plot of calcium activity in yolk sac surface cells at 36 hpf over a total recording time of ∼500 s, with each ROI representing a single cell. (**E**), Representative confocal images of calcium transients on the yolk sac surface of Tg(elavl3:GCaMP6f) embryos at 48 hpf. White arrows point to cells displaying robust calcium flashes. Green fluorescence indicates pan-neuronal Ca2+ dynamics. Time-lapse imaging was acquired using a 30× silicone immersion objective at 6.69 s per frame. Scale bar = 50 μm. (**F**), Statistical raster diagram of calcium events in yolk sac surface cells at 48 hpf across ∼500 s continuous recording; each ROI denotes an individual cell. (**G**), Representative confocal views of calcium flashes on the yolk sac surface of Tg(elavl3:GCaMP6f) zebrafish embryos at 52 hpf. White arrows highlight cells with obvious calcium transients. Green fluorescence reports pan-neuronal Ca2+ signals. Imaging was conducted with a 30× silicone immersion objective at a frame interval of 5.38 s. Scale bar = 50 μm. (**H**), Raster plot profiling calcium activity of yolk sac surface cells at 52 hpf over a total recording period of ∼500 s; each ROI corresponds to one individual cell. (**I**), Quantitative statistics of calcium transient frequency in yolk sac surface cells across different developmental stages. (**J**), Calcium flicker amplitude statistics of yolk sac surface cells in zebrafish embryos across developmental time points. (**K**), Line graph of calcium transient frequency at different developmental stages in zebrafish embryos. Black line, frequency in yolk sac surface cells; red line, frequency in neuronal somata of the brain. For all statistical plots, unpaired Student’s t-test (n > 3 experiments, data are mean ± SD) was conducted for all data with **** P < 0.0001, *** P < 0.001, ** P < 0.1, * P <0.05, ns P ≥ 0.05. The error bars are the standard deviation of the data.

Because elavl3 (also known as HuC) is generally considered a specific marker for mature neurons (Kim et al., 1996; Park et al., 2000), and the traditional view holds that neural cells are absent on the YS surface, we conducted a rigorous analysis of the cellular origin of the observed calcium flickers. Crucially, we do not interpret these GCaMP6f-positive surface cells as bona fide neurons or neural somata residing on the YS. Instead, because our experiments utilized transgenic lines driven by the elavl3 promoter, the most plausible explanation for these transient calcium flashes is that YS surface cells express low, basal levels of elavl3 during early embryonic development. Previous literature has demonstrated that elavl3 promoter activity is not strictly confined to mature neurons and can display transient expression in non-neural ectodermal lineages and sensory primordia. For example, Park et al. (Park et al., 2000) found that elavl3 is expressed in early photoreceptor precursor cells in the developing zebrafish retina, and these cells downregulate elavl3 upon maturation. Thanks to the high sensitivity of GCaMP6f, even low-level promoter-driven expression in these surface cells is sufficient to yield detectable calcium transients. Therefore, these collective calcium flashes reflect primitive intercellular coordination among YS surface cells responding to microenvironmental cues, here the innervation of neural axons, rather than intrinsic firing of YS-resident neurons.

Calcium signaling, the most ubiquitous intracellular second messenger, plays a central regulatory role in cell proliferation, migration, polarity establishment, and morphogenesis (Webb and Miller, 2000; Slusarski and Pelegri, 2007; Brodskiy and Zartman, 2018). It is known that in the nervous system, spontaneous calcium activity within neurons and neuronal growth cones is essential for axon pathfinding, neural network construction, and circuit maturation (Plazas et al., 2013; Webb and Miller, 2013; Yuen et al., 2013). Being innervated by the central neural system, the YS needs to sense internal and external chemical changes. Elavl3-positive cells may act as chemoreceptors, transmitting environmental information to other embryonic organ systems via calcium signals. To test these possibilities, future studies will need to employ single-cell RNA sequencing to determine the transcriptomic profiles of these cells, and immunohistochemistry to detect whether they express additional neuronal or stem cell markers. Moreover, manipulating the activity of these cells using optogenetic or chemogenetic techniques and observing the effects on embryonic development and YS function will help reveal their specific biological roles. Regardless of the cellular origin, the population calcium burst activity exhibited by these cells indicates the existence of coordinated functional connections among them.

### Neuronal Pruning and Metabolic Stress-Induced Lipid Droplet Crystallization

During long-term imaging, we observed dynamic remodeling of the neural network on the YS surface. We found that some nerve fibers suddenly exhibited local calcium overload, manifested as a sharp increase in GCaMP6f fluorescence intensity, after which these fibers gradually fragmented and disappeared within minutes (**Fig.3A and B; Movie S3)**. This phenomenon closely resembles the widespread neuronal pruning or local degeneration observed during neural system development, representing an important hallmark of network refinement and functional optimization (Wang et al., 2026). We speculate that this calcium-dependent pruning mechanism can eliminate redundant, misconnected, or dysfunctional fibers, thereby ensuring the precision and efficacy of the YS surface neural network (Hua et al., 2005; Kanamori et al., 2013).

**Fig. 3.**
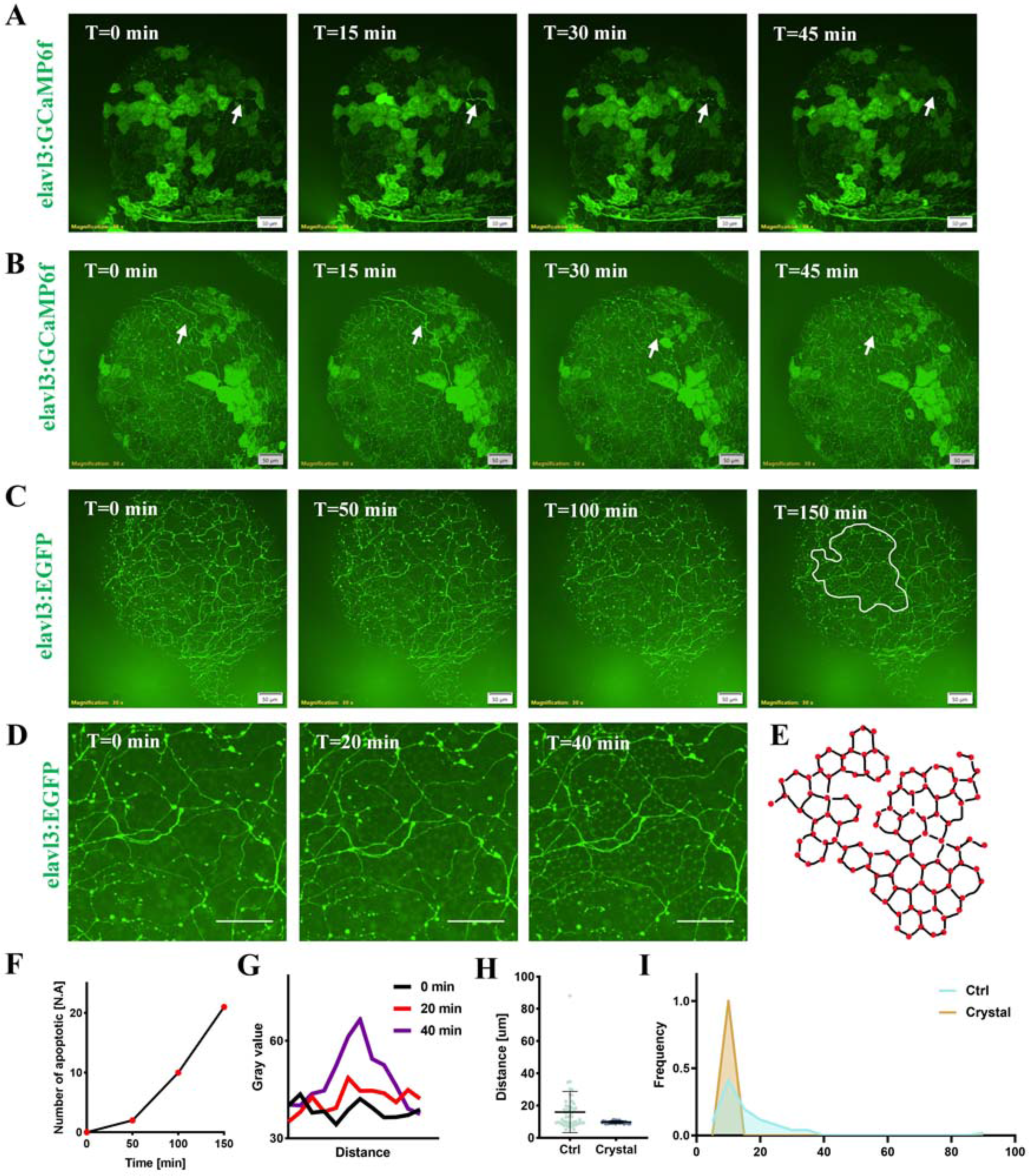
Activity-dependent neuronal pruning and stress-induced lipid droplet crystallization in the yolk sac. (**A, B**), Representative confocal time-lapse images illustrating activity-dependent structural refinement of neural networks on the yolk sac surface of Tg(elavl3:GCaMP6f) zebrafish embryos. White arrows denote the refined neural structures. Green fluorescence signals label pan-neuronal calcium activity. Time-lapse imaging was acquired using a 30× silicone immersion objective with an interval of 15 min. Scale bar = 50 μm. (**C**), Representative confocal micrographs showing crystalline aggregation of lipid droplets within the yolk sac of moribund Tg(elavl3:EGFP) zebrafish embryos. White labeled regions indicate areas with crystalline structures, and green fluorescence labels pan-neuronal signals. Imaging was performed with a 30× silicone immersion objective at a 10 min frame interval. Scale bar = 50 μm. (**D**), Representative magnified confocal images showing lipid droplets in the yolk sac of Tg(elavl3:EGFP) zebrafish embryos before and after crystalline aggregation. Scale bar = 50 μm. (**E**), Schematic diagram of the structure formed by crystalline aggregation of lipid droplets. (**F**), Quantification of apoptotic axon numbers at different time points, based on the live-cell imaging data shown in (**C**). (**G**), Temporal changes in gray value measured at the same location on the yolk sac surface during lipid droplet crystallization, derived from the live-cell imaging data used to generate the schematic in (**E**). (**H**), Statistical comparison of the nearest-neighbor distance between lipid droplets under normal (non-crystalline) and stress-induced crystalline conditions in the yolk sac. (**I**), Frequency distribution analysis of inter-droplet distances for lipid droplets in non-crystalline and crystalline yolk sac states. For all statistical plots (n > 3 experiments) was conducted for all data. The error bars are the standard deviation of the data.

Furthermore, we observed that at later imaging stages, when embryos were approaching death due to phototoxicity from laser exposure or other stress factors, the lipid droplets (LDs) within the YS underwent significant morphological and distributional changes (**Fig.3C-E; Movie S4)**. It should be noted that no specific lipophilic dyes were used in these experiments. Nevertheless, individual lipid droplets were clearly distinguishable under confocal and bright-field imaging. This optical visualization is primarily attributed to the significant refractive index mismatch between the lipid-rich core of the droplets and the surrounding aqueous cytoplasmic environment, which creates high optical contrast, along with weak background signal scattering. Under stress conditions, these optically distinct lipid droplets shifted from a randomly dispersed state to a highly ordered, hexagonal-like aggregated configuration (**Fig. 3C-E**). The observed lipid droplet crystallization in the yolk sac was accompanied by the collapse of the neural network on the yolk sac surface (**Fig. 3F**). Notably, prominent aggregation signals were detected within approximately 40 min after the onset of crystallization (**Fig. 3G**). In normally developing embryos, lipid droplets were dispersed within the YS, with an average nearest neighbor distance (NND) of approximately 15 μm between adjacent droplets. In contrast, when embryos were dying, lipid droplets rapidly aggregated into regular crystalline-like structures; the average NND decreased markedly to 8 μm, and the size and shape of the droplets became more uniform (**Fig.3H and I)**.

These results indicate that the ordered arrangement and dispersed state of lipid droplets are important hallmarks of YS internal environment homeostasis, whereas crystalline aggregation of lipid droplets reflects loss of YS membrane integrity and collapse of the intracellular milieu e.g., pH, ion concentration (Shimobayashi and Ohsaki, 2019; Zadoorian et al., 2023). Lipid droplets are not only organelles for lipid storage but also important platforms for intracellular signaling and stress responses. Under normal physiological conditions, lipid droplets are dispersed to ensure efficient lipid metabolism and transport. When cells are subjected to stress or homeostatic disruption, lipid droplets aggregate; this may be a cytoprotective mechanism that reduces the risk of lipid peroxidation by decreasing droplet surface area, or enhances stress resistance by recruiting relevant stress proteins. However, when stress exceeds the tolerance limit, lipid droplet aggregation becomes irreversible, ultimately leading to cell death. Therefore, the distribution state of lipid droplets can serve as a sensitive, non-invasive biosensor for assessing embryonic health and stress levels.

### Pre-vascular Directional Blood Flow on the Yolk Sac Surface

To study blood flow on the YS surface and its relationship with vascular development, we used the transgenic line Tg(elavl3:EGFP; fli-1:mCherry), which specifically expresses green fluorescent protein EGFP in pan-neuronal cells and red fluorescent protein mCherry in vascular endothelial cells (Park et al., 2000; Lawson and Weinstein, 2002). We performed bright-field and fluorescence imaging on the same embryo to simultaneously observe blood cell movement and the distribution of vascular endothelial cells.

Fluorescence imaging showed that at 30 hpf, fli1-positive vascular endothelial cells began to sprout and grow onto the yolk sac surface; at this time, the endothelial cells on the YS surface were still scattered and isolated, without forming continuous lumens (**Fig.4A; Movie S5**). Between 30 and 42 hpf, endothelial cells on the yolk sac surface formed vascular channels, representing an important time window; they adopted a flattened channel-like structure, which is markedly different from the lumen-like structures formed in the brain. However, surprisingly, bright-field imaging revealed that around 30 hpf, before vascular endothelial cells had formed channels on the yolk sac surface, blood cells could already be observed entering the YS surface and undergoing directional flow (**Fig.4B; Movie S5)**. We quantified the blood cell flow velocity, and the results showed an average velocity of approximately 200 μm/min (**Fig.4C)**, consistent with previously reported early blood circulation velocities in zebrafish embryos.

**Fig. 4.**
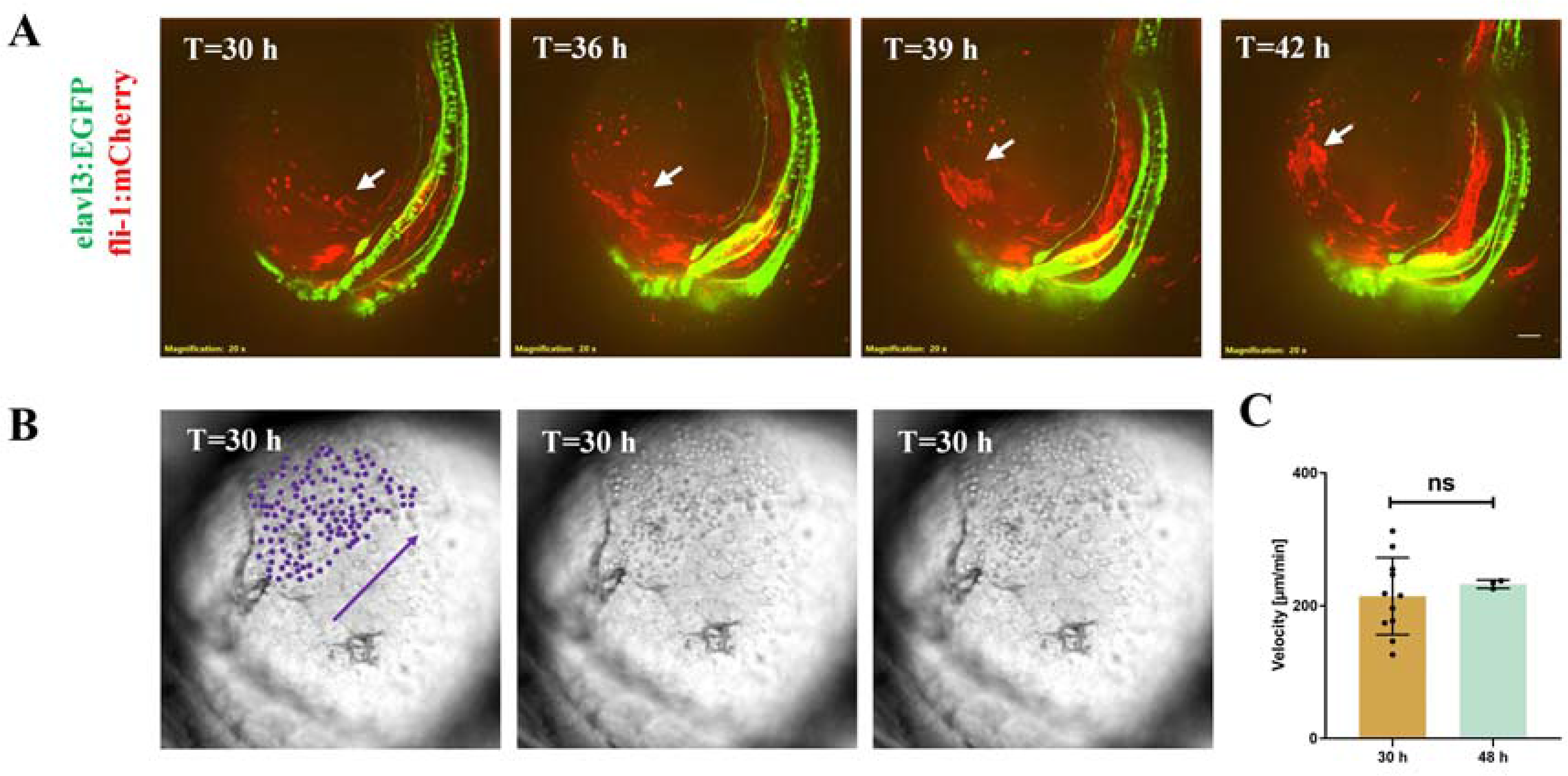
Pre-vascular directional blood flow on the yolk sac surface prior to endothelial tube formation. (**A**), Representative confocal time-lapse images of neural and vascular patterning around the yolk sac in double-transgenic Tg(elavl3:EGFP; fli-1:mCherry) zebrafish embryos starting at 30 hpf. Green fluorescence labels pan-neuronal structures, and red fluorescence marks vascular endothelial cells. White arrows indicate endothelial signals on the yolk sac surface. Imaging was performed using a 20× air objective with a time interval of 30 min. Scale bar = 50 μm. (**B**), Representative brightfield micrograph showing blood circulation on the yolk sac surface of zebrafish embryos at 30 hpf. Purple dots denote circulating blood cells, and arrows indicate the directional route of blood transport. (**C**), Quantitative statistical plot of blood cell flow velocity on the yolk sac surface across different developmental stages of zebrafish embryos. For all statistical plots (n > 3 experiments) was conducted for all data. For all statistical plots, unpaired Student’s t-test (n > 3 experiments, data are mean ±SD) was conducted for all data with **** P < 0.0001, *** P < 0.001, ** P < 0.1, * P <0.05, ns P ≥ 0.05. The error bars are the standard deviation of the data.

This result clearly demonstrates the existence of a blood transport route on the YS surface independent of vascular endothelial cells in early zebrafish development. Our previous work demonstrated that three distinct circulatory loops are established during early zebrafish embryogenesis: one confined to the brain, one spanning the trunk, and a third localized entirely on the surface of the yolk sac. Specifically, this vitelline circulation exhibits a well-organized directional flow, where blood exits the heart to course across one side of the yolk sac, before draining back into the heart from the contralateral side (Wang et al., 2026). We speculate that this route may be an interstitial flow channel formed by the extracellular matrix on the YSL surface, or a temporary conduit created by membrane invaginations of the YSL itself. This “flow before tube” phenomenon has important biomechanical significance (Xu and Cleaver, 2011; Goetz et al., 2014). The traditional view holds that blood circulation in vertebrates must rely on vascular endothelial cells fully assembling into closed tubes (Jin et al., 2005; Potente and Mäkinen, 2017). However, recent intravital imaging studies have revealed that, before endothelial cells complete definitive lumen formation, a nascent plasma flow driven by cardiac pumping already achieves directional flow within loose intercellular spaces (Kamei et al., 2006; Herbert et al., 2009).

We speculate that the fluid shear stress generated by this “tubeless directional flow” can activate mechanosensitive channels (such as Piezo1) and primary cilia on endothelial cells, and through mechanotransduction, inversely catalyze intracellular vacuole fusion and lumenization in endothelial cells, thereby dynamically shaping and pruning the primitive vascular network (Wang et al., 2010; Delling et al., 2016; Gebala et al., 2016; Liu et al., 2020). This flow-guided angiogenesis mechanism ensures that the formation of the vascular system matches the demands of blood circulation, representing an efficient developmental regulatory strategy. In addition, non-vascular-dependent blood flow may play an important role in oxygen and nutrient transport. Before the vascular system is fully formed, the metabolic demand of the embryo is relatively low, and substance transport via interstitial flow is sufficient to meet the embryo’s needs. As development proceeds and metabolic demand increases, the vascular system gradually forms and takes over the transport function. This transition from non-vascular to vascular transport represents an important evolutionary adaptation in vertebrate embryonic development.

## Discussion

In this study, using multiple transgenic zebrafish lines and long-term high-resolution live imaging, we have systematically revealed for the first time the dynamic processes of neural development, calcium activity, hemodynamics, and metabolic homeostasis on the YS surface of zebrafish embryos. Our findings challenge the traditional view of YS function, demonstrating that, with neural innervation, the YS functions beyond a nutrient storage organ or biochemical regulator. It also serves as a development hub, with complex synergistic interactions among the neural, vascular, and metabolic systems on its surface.

We find that during zebrafish development, neuronal growth cones cover the YS surface, forming a complex neural network from cranial neuron projections. Although YS structures were thought to degrade upon absorption, we propose that, innervated and directed by neurons, yolk executes transient biological function and is –reused: nerve fibers reorganize it into a primitive scaffold for the autonomic nervous system. We also observe elavl3□cells on the YS with calcium bursts, a hallmark of neural circuit establishment. Upon embryo death, lipid droplets crystallize, indicating their role in homeostasis and as a biosensor for health. Finally, we discover “flow before tube”: directional blood flow on the YS prior to endothelial tube formation provides oxygen/nutrients and shear stress to guide endothelial migration and tube formation. Together, these findings reveal the YS as a dynamic hub for neurodevelopment, chemosensation, vascular patterning, and stress sensing, far beyond nutrient storage.

This study is the first systematic investigation of neural, vascular, and metabolic dynamics on the zebrafish YS surface. However, key questions remain. First, the specific functions of the brain-YS circuit and the microscopic synaptic structure between neuron axons and YS surface cells are yet to be elucidated. Using optogenetic techniques to activate or inhibit these neurons and observing the subsequent effects on embryonic development, YS metabolism, and visceral organ function will be a key future direction. Second, the precise structural and molecular mechanisms of non-vascular blood flow are not determined. Overall, our results uncover the YS as a developmental regulator, offering a new perspective on the crosstalk among neural, vascular, and metabolic systems in early vertebrate development, and have implications for diagnosing and treating embryonic developmental disorders.

## Acknowledgments

We thank Peishi Wang and Mingyang Chen for helping maintain the transgenetic zebrafish lines; Runjie Yu from Oujiang Laboratory for assistance in spinning disk confocal microscopy imaging, respectively; and Deepseek v2.2.1 for text polishing. This study was supported by taxpayers of China through National Natural Science Foundation of China (Distinguished Young Scholars Funding, Overseas; Young Scientists Fund - Category C) and Wuhan University (Talents Startup Funding).

## Author contributions

B.L. conceptualized the project, B.L. Z.D.W. designed the research, Z.D.W and L.T. performed the experiments, Z.D.W. analyzed the data. All authors discussed the results and wrote the paper.

## Data availability

The data and analysis code that support the findings of this study are available upon reasonable request to B.L..

## References

1. Avitan L, Pujic Z, Mölter J, Zhu S, Sun B, Goodhill GJ (2021) Spontaneous and evoked activity patterns diverge over development. Elife 10.

2. Bader HL, Keene DR, Charvet B, Veit G, Driever W, Koch M, Ruggiero F (2009) Zebrafish collagen XII is present in embryonic connective tissue sheaths (fascia) and basement membranes. Matrix Biol 28:32–43.

3. Bischof J, Driever W (2004) Regulation of hhex expression in the yolk syncytial layer, the potential Nieuwkoop center homolog in zebrafish. Dev Biol 276:552–562.

4. Brodskiy PA, Zartman JJ (2018) Calcium as a signal integrator in developing epithelial tissues. Phys Biol 15:051001.

5. Carvalho L, Heisenberg CP (2010) The yolk syncytial layer in early zebrafish development. Trends Cell Biol 20:586–592.

6. Chen Z, He M, Wang H, Li X, Qin R, Ye D, Zhai X, Zhu J, Zhang Q, Hu P, Shui G, Sun Y (2024) Intestinal DHA-PA-PG axis promotes digestive organ expansion by mediating usage of maternally deposited yolk lipids. Nat Commun 15:9769.

7. Chia K, Klingseisen A, Sieger D, Priller J (2022) Zebrafish as a model organism for neurodegenerative disease. Front Mol Neurosci 15:940484.

8. Delling M, Indzhykulian AA, Liu X, Li Y, Xie T, Corey DP, Clapham DE (2016) Primary cilia are not calcium-responsive mechanosensors. Nature 531:656–660.

9. Diez-Pinel G, Muratore A, Ruhrberg C, Canu G (2026) Discovery of New Markers for Haemogenic Endothelium and Haematopoietic Progenitors in the Mouse Yolk Sac. J Dev Biol 14.

10. Dunn TW, Mu Y, Narayan S, Randlett O, Naumann EA, Yang CT, Schier AF, Freeman J, Engert F, Ahrens MB (2016) Brain-wide mapping of neural activity controlling zebrafish exploratory locomotion. Elife 5:e12741.

11. Feldman B, Gates MA, Egan ES, Dougan ST, Rennebeck G, Sirotkin HI, Schier AF, Talbot WS (1998) Zebrafish organizer development and germ-layer formation require nodal-related signals. Nature 395:181–185.

12. Fukazawa C, Santiago C, Park KM, Deery WJ, Gomez de la Torre Canny S, Holterhoff CK, Wagner DS (2010) poky/chuk/ikk1 is required for differentiation of the zebrafish embryonic epidermis. Dev Biol 346:272–283.

13. Furukawa F, Aoyagi A, Sano K, Sameshima K, Goto M, Tseng YC, Ikeda D, Lin CC, Uchida K, Okumura SI, Yasumoto K, Jimbo M, Hwang PP (2024) Gluconeogenesis in the extraembryonic yolk syncytial layer of the zebrafish embryo. PNAS Nexus 3:pgae125.

14. Gebala V, Collins R, Geudens I, Phng LK, Gerhardt H (2016) Blood flow drives lumen formation by inverse membrane blebbing during angiogenesis in vivo. Nat Cell Biol 18:443–450.

15. Goetz JG, Steed E, Ferreira RR, Roth S, Ramspacher C, Boselli F, Charvin G, Liebling M, Wyart C, Schwab Y, Vermot J (2014) Endothelial cilia mediate low flow sensing during zebrafish vascular development. Cell Rep 6:799–808.

16. Herbert SP, Huisken J, Kim TN, Feldman ME, Houseman BT, Wang RA, Shokat KM, Stainier DY (2009) Arterial-venous segregation by selective cell sprouting: an alternative mode of blood vessel formation. Science 326:294–298.

17. Hua JY, Smear MC, Baier H, Smith SJ (2005) Regulation of axon growth in vivo by activity-based competition. Nature 434:1022–1026.

18. Iturri L, Freyer L, Biton A, Dardenne P, Lallemand Y, Gomez Perdiguero E (2021) Megakaryocyte production is sustained by direct differentiation from erythromyeloid progenitors in the yolk sac until midgestation. Immunity 54:1433–1446.e1435.

19. Jin SW, Beis D, Mitchell T, Chen JN, Stainier DY (2005) Cellular and molecular analyses of vascular tube and lumen formation in zebrafish. Development 132:5199–5209.

20. Kamei M, Saunders WB, Bayless KJ, Dye L, Davis GE, Weinstein BM (2006) Endothelial tubes assemble from intracellular vacuoles in vivo. Nature 442:453–456.

21. Kanamori T, Kanai MI, Dairyo Y, Yasunaga K, Morikawa RK, Emoto K (2013) Compartmentalized calcium transients trigger dendrite pruning in Drosophila sensory neurons. Science 340:1475–1478.

22. Kawahara A, Nishi T, Hisano Y, Fukui H, Yamaguchi A, Mochizuki N (2009) The sphingolipid transporter spns2 functions in migration of zebrafish myocardial precursors. Science 323:524–527.

23. Kiener TK, Selptsova-Friedrich I, Hunziker W (2008) Tjp3/zo-3 is critical for epidermal barrier function in zebrafish embryos. Dev Biol 316:36–49.

24. Kim CH, Ueshima E, Muraoka O, Tanaka H, Yeo SY, Huh TL, Miki N (1996) Zebrafish elav/HuC homologue as a very early neuronal marker. Neurosci Lett 216:109–112.

25. Kimmel CB, Warga RM, Schilling TF (1990) Origin and organization of the zebrafish fate map. Development 108:581–594.

26. Lawson ND, Weinstein BM (2002) In vivo imaging of embryonic vascular development using transgenic zebrafish. Dev Biol 248:307–318.

27. Lessman CA (2011) The developing zebrafish (Danio rerio): a vertebrate model for high-throughput screening of chemical libraries. Birth Defects Res C Embryo Today 93:268–280.

28. Liu TT, Du XF, Zhang BB, Zi HX, Yan Y, Yin JA, Hou H, Gu SY, Chen Q, Du JL (2020) Piezo1-Mediated Ca(2+) Activities Regulate Brain Vascular Pathfinding during Development. Neuron 108:180–192.e185.

29. Park HC, Kim CH, Bae YK, Yeo SY, Kim SH, Hong SK, Shin J, Yoo KW, Hibi M, Hirano T, Miki N, Chitnis AB, Huh TL (2000) Analysis of upstream elements in the HuC promoter leads to the establishment of transgenic zebrafish with fluorescent neurons. Dev Biol 227:279–293.

30. Plazas PV, Nicol X, Spitzer NC (2013) Activity-dependent competition regulates motor neuron axon pathfinding via PlexinA3. Proc Natl Acad Sci U S A 110:1524–1529.

31. Potente M, Mäkinen T (2017) Vascular heterogeneity and specialization in development and disease. Nat Rev Mol Cell Biol 18:477–494.

32. Quinlivan VH, Farber SA (2017) Lipid Uptake, Metabolism, and Transport in the Larval Zebrafish. Front Endocrinol (Lausanne) 8:319.

33. Rosch RE, Burrows DRW, Lynn CW, Ashourvan A (2024) Spontaneous Brain Activity Emerges from Pairwise Interactions in the Larval Zebrafish Brain. Phys Rev X 14.

34. Sakaguchi T, Kikuchi Y, Kuroiwa A, Takeda H, Stainier DY (2006) The yolk syncytial layer regulates myocardial migration by influencing extracellular matrix assembly in zebrafish. Development 133:4063–4072.

35. Sampedro MF, Izaguirre MF, Sigot V (2018) E-cadherin expression pattern during zebrafish embryonic epidermis development. F1000Res 7:1489.

36. Shimobayashi SF, Ohsaki Y (2019) Universal phase behaviors of intracellular lipid droplets. Proc Natl Acad Sci U S A 116:25440–25445.

37. Slusarski DC, Pelegri F (2007) Calcium signaling in vertebrate embryonic patterning and morphogenesis. Dev Biol 307:1–13.

38. Sun T, Tao Y, Zeng Y, Ai R, Li T, Li X, Li X, He R, Jia Z, Sun X, Feng Z, Liu X, Kong X, Huang L, Chen L, Ni J, Chen L, Dai W (2026) Increased yolk lipid mobilization promotes zebrafish post-segmentation growth via an Hnf4-lipoprotein axis. Cell Rep 45:117295.

39. Takeuchi M, Yamaguchi S, Yonemura S, Kakiguchi K, Sato Y, Higashiyama T, Shimizu T, Hibi M (2015) Type IV Collagen Controls the Axogenesis of Cerebellar Granule Cells by Regulating Basement Membrane Integrity in Zebrafish. PLoS Genet 11:e1005587.

40. Wang Y, Kaiser MS, Larson JD, Nasevicius A, Clark KJ, Wadman SA, Roberg-Perez SE, Ekker SC, Hackett PB, McGrail M, Essner JJ (2010) Moesin1 and Ve-cadherin are required in endothelial cells during in vivo tubulogenesis. Development 137:3119–3128.

41. Wang Z, Tian L, Li B (2026) Neurogenesis Leads Early Development in Zebrafish. bioRxiv:2025.2011.2012.687769.

42. Webb SE, Miller AL (2000) Calcium signalling during zebrafish embryonic development. Bioessays 22:113–123.

43. Webb SE, Miller AL (2013) Calcium signaling in extraembryonic domains during early teleost development. Int Rev Cell Mol Biol 304:369–418.

44. Xu K, Cleaver O (2011) Tubulogenesis during blood vessel formation. Semin Cell Dev Biol 22:993–1004.

45. Yuen MY, Webb SE, Chan CM, Thisse B, Thisse C, Miller AL (2013) Characterization of Ca(2+) signaling in the external yolk syncytial layer during the late blastula and early gastrula periods of zebrafish development. Biochim Biophys Acta 1833:1641–1656.

46. Zadoorian A, Du X, Yang H (2023) Lipid droplet biogenesis and functions in health and disease. Nat Rev Endocrinol 19:443–459.

47. Zhao A, Gruntman E, Nern A, Iyer N, Rogers EM, Koskela S, Siwanowicz I, Dreher M, Flynn MA, Laughland C, Ludwig H, Thomson A, Moran C, Gezahegn B, Bock DD, Reiser MB (2025) Eye structure shapes neuron function in Drosophila motion vision. Nature 646:135–142.

