## Supplementary Information for "Neural Innervation Invigorates Yolk Sac Biological Functions beyond Nutrient Reservoir during Zebrafish Embryo Development"

Zhengduo Wang^1^, Li Tian^2^ and Bo Li^1, *^

1. The Institute for Advanced Studies, Wuhan University, Wuhan 430072, P. R. China.
2. Department of Physics, Wenzhou University, Wenzhou 325000, P. R. China.

**This PDF file includes:**

Legends for Movies S1 to S5

**Other supplementary materials for this manuscript include the following:**

Movies S1 to S5

**Legends for Supplementary Movies**

**Movie S1.** **Wide-field time-lapse confocal imaging was performed to capture the development of the neural network on the yolk sac surface of zebrafish (elavl3:EGFP) embryos.** A 30x silicone oil immersion objective lens was used. All neuronal structures were labeled with EGFP (excitation at 488 nm; laser power 20%; exposure time 200 ms). Imaging started at 24 hours post‑fertilization and was conducted every 10 minutes for 14 hours. Scale bar = 50 μm.

**Movie S2.** **Wide-field time-lapse confocal imaging of calcium transients on the yolk sac surface of zebrafish (elavl3:GCaMP6f) embryos.** Four time points (24, 36, 48, and 52 hours post‑fertilization) are shown from left to right. A 30x silicone oil immersion objective lens was used. Calcium signals were labeled with GCaMP6f (excitation at 488 nm; laser power 20%; exposure time 200 ms). Imaging intervals: 24 hpf-every 10.42 s; 36 hpf-every 5.58 s; 48 hpf-every 6.69 s; 52 hpf-every 5.38 s. Scale bar = 50 μm

**Movie S3.** **Wide-field time-lapse confocal imaging was performed to visualize the refinement of local neural network structures on the yolk sac surface of zebrafish (elavl3:GCaMP6f) embryos at 40 hours post‑fertilization.** A 30× silicone oil immersion objective lens was used. Calcium signals were labeled with GCaMP6f (excitation at 488 nm; laser power 20%; exposure time 200 ms). Images were acquired every 15 minutes for a total duration of approximately 12 hours. Scale bar = 50μm.

**Movie S4.** **Wide-field time-lapse confocal imaging was performed to visualize lipid droplet crystallization on the yolk sac surface of zebrafish (elavl3:EGFP) embryos at 36 hours post‑fertilization.** A 30× silicone oil immersion objective lens was used. Neurons were labeled with EGFP (excitation at 488 nm; laser power 20%; exposure time 200 ms). Images were acquired every 10 minutes for a total duration of approximately 3.5 hours. Scale bar = 50μm.

**Movie S5.** **Time-lapse confocal imaging of vascular/neural network formation and blood cell circulation on the yolk sac surface of zebrafish embryos.** A 20× objective lens was used for imaging. Vascular endothelial cells were labeled with mCherry (excitation at 561 nm; laser power 20%; exposure time 200 ms), and neurons were labeled with EGFP (excitation at 488 nm; laser power 20%; exposure time 200 ms). Images were acquired every 20 minutes for a total duration of approximately 20 hours. Scale bar = 50 μm. Left: Wide-field time-lapse confocal imaging at 24 hours post‑fertilization. A 20× objective lens was used. Vascular endothelial cells were labeled with mCherry (excitation 561 nm; laser power 20%; exposure 200 ms), and neurons with EGFP (excitation 488 nm; laser power 20%; exposure 200 ms). Images were acquired every 20 minutes for a total duration of approximately 20 hours. Scale bar = 50 μm. Right: Time-lapse confocal imaging of blood cell circulation on the yolk sac surface. A 30× silicone oil‑immersion objective was used. Imaging started at 30 hours post‑fertilization with a 1‑second interval.
